# SARS-CoV-2 nucleocapsid protein dually regulates innate immune responses

**DOI:** 10.1101/2021.02.17.431755

**Authors:** Yinghua Zhao, Liyan Sui, Ping Wu, Wenfang Wang, Guangyun Tan, Zedong Wang, Yang Yu, Zhijun Hou, Guoqing Wang, Quan Liu

**Author notes:** Correspondence to: Guoqing Wang, Quan Liu.

## Abstract

The recently emerged severe acute respiratory syndrome coronavirus 2 (SARS-CoV-2), the causative agent of the ongoing global pandemic of COVID-19, may trigger immunosuppression in the early stage and a cytokine storm in the late stage of infection, however, the underlying mechanisms are not well understood. Here we demonstrated that the SARS-CoV-2 nucleocapsid (N) protein dually regulated innate immune responses, i.e., the low-dose N protein suppressed type I interferon (IFN-I) signaling and inflammatory cytokines, whereas high-dose N protein promoted IFN-I signaling and inflammatory cytokines. Mechanistically, the SARS-CoV-2 N protein interacted with the tripartite motif protein 25 (TRIM25), thereby dually regulating the phosphorylation and nuclear translocation of IRF3, STAT1 and STAT2. Additionally, low-dose N protein combined with TRIM25 could suppress retinoic acid-inducible gene I (RIG-I) ubiquitination and activation. Our findings revealed a regulatory mechanism of innate immune responses by the SARS-CoV-2 N protein, which would contribute to understanding the pathogenesis of SARS-CoV-2 and other SARS-like coronaviruses, and development of more effective strategies for controlling COVID-19.

## INTRODUCTION

Severe acute respiratory syndrome coronavirus 2 (SARS-CoV-2) is a new member in the *Coronaviridae* family that is genetically related to SARS-CoV and Middle East respiratory syndrome coronavirus (MERS-CoV) (1, 2). The resulting coronavirus disease 2019 (COVID-19) pandemic has caused tens millions of infections worldwide. The elderly, especially those with comorbid lung disease, heart disease, diabetes, or obesity, have the most severe clinical symptoms, leading to a high mortality (3). SARS-CoV-2 triggers an immunosuppression in the early stage of infection, which contributes to uncontrolled coronaviral replication (4–6). This overwhelming coronaviral replication, in turn, stimulates an aberrant unchecked cytokine release, known as “cytokine storm“, leading to severe acute respiratory distress syndrome (ARDS), pneumonia, and multiple organs failure (7). Understanding the underlying mechanisms is conducive to development of more effective strategies for COVID-19.

Type I interferons (IFN-I) play a critical role in the host antiviral process at the early stage of infection, whose production is rapidly triggered by recognition of pathogen-associated molecular patterns (PAMPs) through host pattern recognition receptors (PRRs) (8). Like most RNA viruses, coronaviral RNA is recognized by retinoic acid-inducible gene I (RIG-I) and melanoma differentiation-associated gene 5 (MDA5), which activate various signaling pathways that finally lead to the production of IFN-I, interferon stimulated genes (ISGs), and proinflammatory cytokines (9).

Several proteins of SARS-CoV-2 have the IFN-antagonizing properties to overcome its antiviral responses (10, 11). SARS-CoV-2 nucleocapsid (N) protein is a 419-amino acid structural protein that plays a central role in the viral assembly, and interacts with RIG-I to inhibit IFN-β production (12). Here we demonstrated that the SARS-CoV-2 N protein dually regulated IFN-I production, i.e., the low-dose SARS-CoV-2 N protein suppressed IFN-I production via sequestering TRIM25 to inhibit RIG-I ubiquitination, while high-dose N protein promoted IFN-I signaling and inflammatory cytokine expression. Additionally, SARS-CoV N protein was also found to be able to regulate IFN-I production bidirectionally. Our findings revealed an innate immune regulation mechanism mediated by the SARS-CoV-2 N protein, which would contribute to understanding the pathogenesis of SARS-CoV-2 N and SARS-like coronaviruses.

## RESULTS

### N protein dually regulates IFN-I and inflammatory cytokines production

To explore the effect of SARS-CoV-2 N protein on IFN signaling, we constructed the expression plasmid of SARS-CoV-2 N protein (Fig. S1) and examined its regulation of IFN-I production by dual luciferase reporter experiments, in which SARS-CoV N protein was used as a control. The expression of SARS-CoV-2 and SARS-CoV N proteins were confirmed by Western blot (Fig. 1A, B down). We unexpectedly found that low-dose (0.25 μg) SARS-CoV-2 N protein significantly reduced the promoter activity of IFN-β and interferon-stimulated response elements (ISRE) stimulated by poly(I:C), while high-dose (1 μg) SARS-CoV-2 N protein increased the promoter activity (Fig. 1A, B). Similar phenomenon was also observed in SARS-CoV N transfected cells (Fig. 1A, B). We further confirmed the effects of SARS-CoV-2 and SARS-CoV N proteins on the transcription levels of IFN-I and ISGs. Quantitative real-time PCR (qPCR) showed that low-dose (0.25 μg) N proteins were able to effectively inhibit poly(I:C)-induced *IFNA*, *IFNB1*, *ISG15* (interferon-stimulated gene 15), *MxA* (MX dynamin like GTPase 1) and *OAS1* (2’-5’-oligoadenylate synthetase 1) expression, whereas high-dose (1 μg) N proteins had the contrary effects to promote their expression (Fig. 1C, D, and E). Then, we used HepG2 cells to further verify the dually regulatory effect of SARS-CoV-2 N protein on IFN-I and ISGs production induced by poly(I:C) (Fig. S2A-E). In addition, SARS-CoV-2 N protein in different doses also had a contrary effect in the absence of poly(I:C) (Fig. S2C-E). These results showed that low-dose SARS-CoV-2 and SARS-CoV N proteins suppresses the expression of IFN-I and downstream ISGs, while high-dose N proteins promotes the IFN-I signaling.

**FIG 1.**
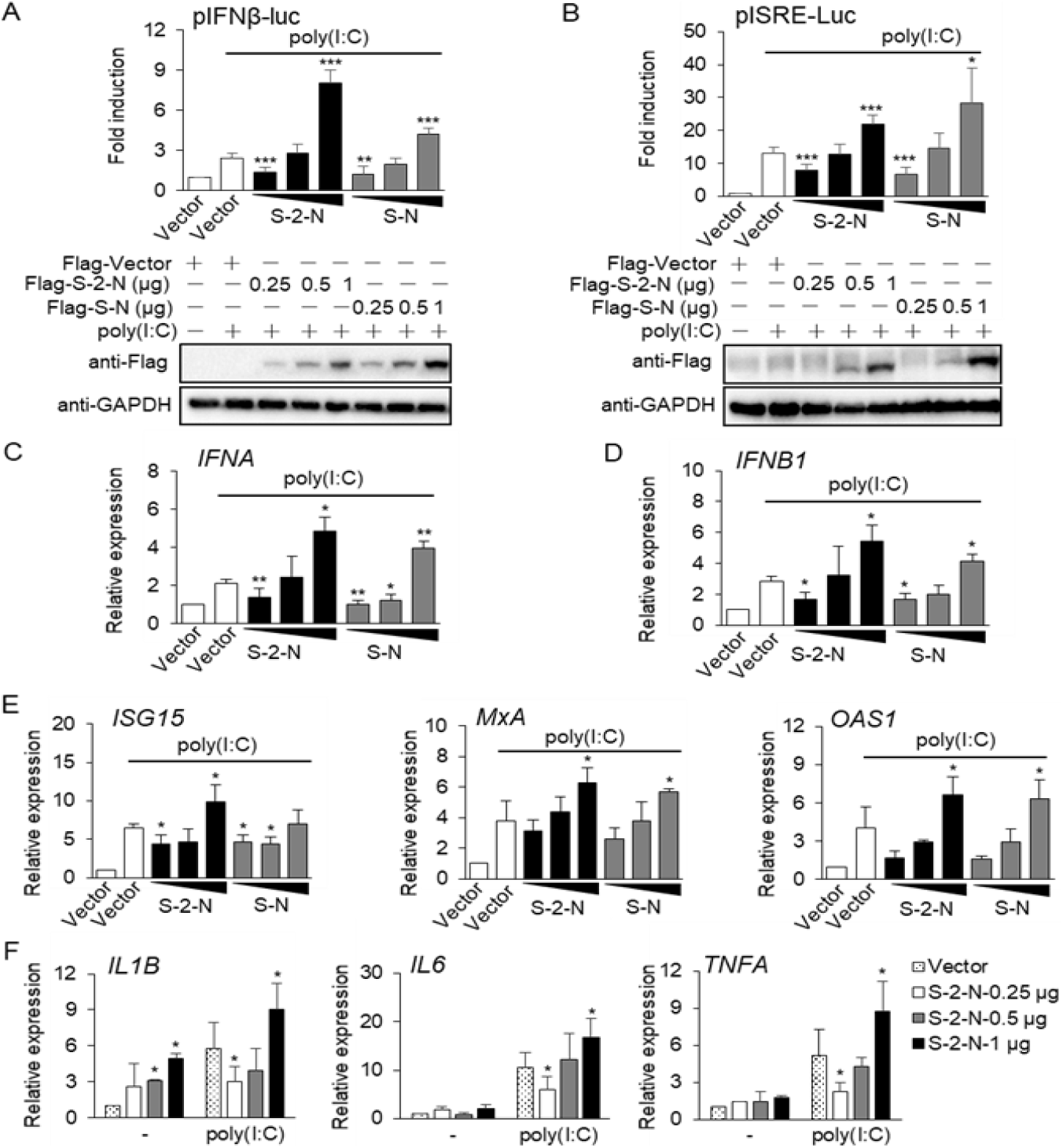
N proteins dually regulate IFN-I and inflammatory cytokines production. (A-B) HEK293T cells were co-transfected with an IFN-β promoter (A) or ISRE reporter plasmid (B), along with control plasmid pGL4.74 and 0.25, 0.5, or 1 μg SARS-CoV-2 or SARS-CoV N proteins (S-2-N or S-N) expressing plasmids, then treated with or without poly(I:C), the empty vector as a control. At 24 h post-transfection (hpt), cells were harvested and luciferase activity was measured (upper); The expression of S-2-N and S-N proteins was detected by Western blot, GAPDH as a loading control (down). (C-E) HEK293T cells were transfected with the 0.25, 0.5, or 1 μg S-2-N or S-N expressing plasmids, then treated with or without poly(I:C), the empty vector as a control. At 24 hpt, the mRNA expression of *IFNA* (C), *IFNB1* (D), and interferon stimulated genes *ISG15*, *OAS1* and *MxA* (E) were examined using qPCR. (F) HEK293T cells were transfected with a dose-gradient S-2-N protein plasmid, then treated with or without poly(I:C). At 24 hpt, the mRNA expression of *IL1B*, *IL6*, and *TNFA* were examined using qPCR. Results shown are the mean ± SD of at least three independent experiments. \**P* < 0.05; \*\**P* < 0.01, and \*\*\**P* < 0.001; two-tailed Student’s *t*-test.

Clinical studies have shown that coronavirus evades innate immunity response during the first ten days of infection, but the subsequent delayed IFN-I response is associated with robust virus replication and severe complications, such as inflammation and “cytokine storm” (5). To determine whether the high-dose SARS-CoV-2 N protein promotes expression of disease-related inflammatory factors, such as interleukin 1 beta (IL1B), interleukin 6 (IL6), and tumor necrosis factor-alpha (TNFA) (13), we transfected HEK293T cells with a dose-gradient SARS-CoV-2 N protein with or without poly(I:C) stimulation; qPCR assays showed that the mRNA levels of *IL1B*, *IL6* and *TNFA* were all increased under the high-dose N protein condition, and reduced under the low-dose condition, whose expression patterns were consistent with that of IFN-I production (Fig. 1F). In addition, we found that SARS-CoV N protein also bidirectionally regulated the expression of *IL1B*, *IL6* and *TNFA* (Fig. S2F). These results indicated that SARS-CoV-2 and SARS-CoV N proteins possess dually regulatory properties of the inflammatory cytokines.

### N protein dually regulates phosphorylation of IRF3, STAT1 and STAT2

The antiviral signal pathway mediated by IFN-I includes two stages: IFN-I production and signal transduction.

In the first stage, viral infection is rapidly sensed by host PRRs, whose signal cascades phosphorylates and dimerizes the transcription factor IRF3 (interferon regulatory factor 3), followed by entering the nucleus to activate the expression of IFN-I (14). In the second stage, secreted IFN-α/β triggers the Janus kinases/signal transducer activator transcription proteins (JAK/STAT) 1 and 2 signaling pathways. Phosphorylated STAT1 and STAT2 can form heterodimers, then combine with IRF9 (interferon regulatory factor 9) to form transcription complex ISGF3 (IFN-stimulated gene factor 3), which translocates to the nucleus to induce IFN-stimulated genes (ISGs), and ultimately elicit an effective antiviral responses (15).

In order to investigate the mechanism of the dually regulated IFN-I signaling by SARS-CoV-2 N protein, we tested the expression and activation of endogenous IRF3, STAT1 and STAT2 upon N protein and poly(I:C) induction. HEK293T cells were transfected with a dose-gradient SARS-CoV-2 N protein, then transfected with or without poly(I:C); SARS-CoV N protein was used as a control. Immunoblot indicated that ectopic expression of SARS-CoV-2 and SARS-CoV N proteins did not affect the total levels of IRF3, STAT1 and STAT2, whereas the phosphorylation of IRF3 (Ser396), STAT1 (Tyr701) and STAT2 (Tyr689) was significantly increased by the N proteins in a dose-dependent manner (Fig. 2). However, p-IRF3, p-STAT1 and p-STAT2 were slightly reduced in the presence of low-dose N proteins compared to the poly(I:C)-activated vector control, probably due to a weak activation of IFN-I signaling by poly(I:C) in the study (Fig. 2), but the decreasing trend was consistent with previous investigations (16, 17). These findings indicated that SARS-CoV-2 and SARS-CoV N proteins dually regulate phosphorylation of IRF3, STAT1, and STAT2.

**FIG 2.**
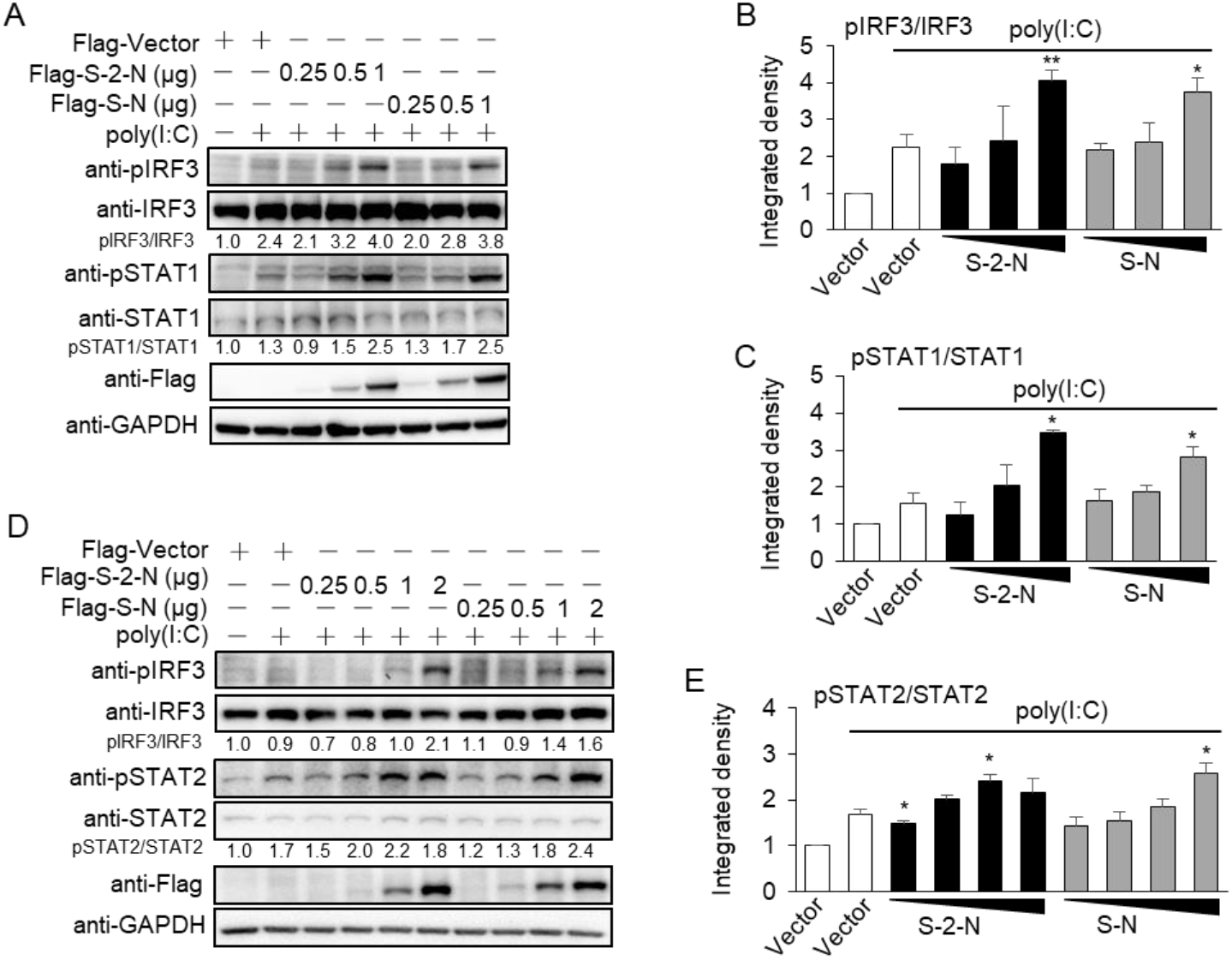
N proteins dually regulate phosphorylation of IRF3, STAT1 and STAT2. (A-C) HEK293T cells were co-transfected with the plasmids expressing Flag-tagged SARS-CoV-2 or SARS-CoV N proteins (S-2-N or S-N) as indicated three increasing doses, then treated with or without poly(I:C), the Flag-vector as a control. At 24 h post-transfection (hpt), the expression of N proteins, the endogenous phosphorylation and total protein levels of IRF3 and STAT1 were detected by Western blot, GAPDH as a loading control (A); the gray-scale statistical analysis of phosphorylation levels of IRF3 (B) and STAT1 (C). (D, E) HEK293T cells were co-transfected with the plasmids expressing Flag-tagged S-2-N or S-N proteins as indicated four increasing doses, then treated with or without poly(I:C), the Flag-vector as a control. At 24 hpt, the expression of N proteins, the endogenous phosphorylation and total protein levels of IRF3 and STAT2 were detected by Western blot, GAPDH as a loading control (D); the gray-scale statistical analysis of phosphorylation level of STAT2 (E). \**P* < 0.05, and \*\**P* < 0.01; two-tailed Student’s *t*-test.

### N protein dually regulates nuclear translocation of IRF3, STAT1 and STAT2

We further determined the effect of different doses of SARS-CoV-2 N protein on nuclear transfer of IRF3, STAT1, and STAT2. We co-transfected IRF3 and high-dose (1 μg) SARS-CoV-2 N protein expression plasmid for 24 h; SARS-CoV N protein was used as a control. Immunofluorescence showed that high-dose N proteins promoted IRF3 to enter the nucleus (Fig. 3A). In contrast, we also co-transfected IRF3 and low-dose (0.25 μg) N proteins plasmids for 24 h, and transfected poly(I:C) to activate IFN signal; Immunofluorescence indicated that low-dose N proteins significantly inhibited poly(I:C) induced nuclear translocation of IRF3 (Fig. 3B). Then we tested the co-localization and nuclear transfer of STAT1 and STAT2 in the presence of different doses of N proteins. SARS-CoV-2 and SARS-CoV N proteins were co-localized with STAT1 and STAT2 in the cytoplasm (Fig. S3). The nuclear transfer of STAT1 and STAT2 were significantly enhanced by high-dose (1 μg) N proteins without poly(I:C) stimulation (Fig. 3C, E), whereas they were inhibited by low-dose (0.25 μg) N proteins in the presence of poly(I:C) stimulation (Fig. 3D, F). These findings indicated that both SARS-CoV-2 and SARS-CoV N proteins dually regulate the nuclear translocation of IRF3, STAT1 and STAT2.

**FIG 3.**
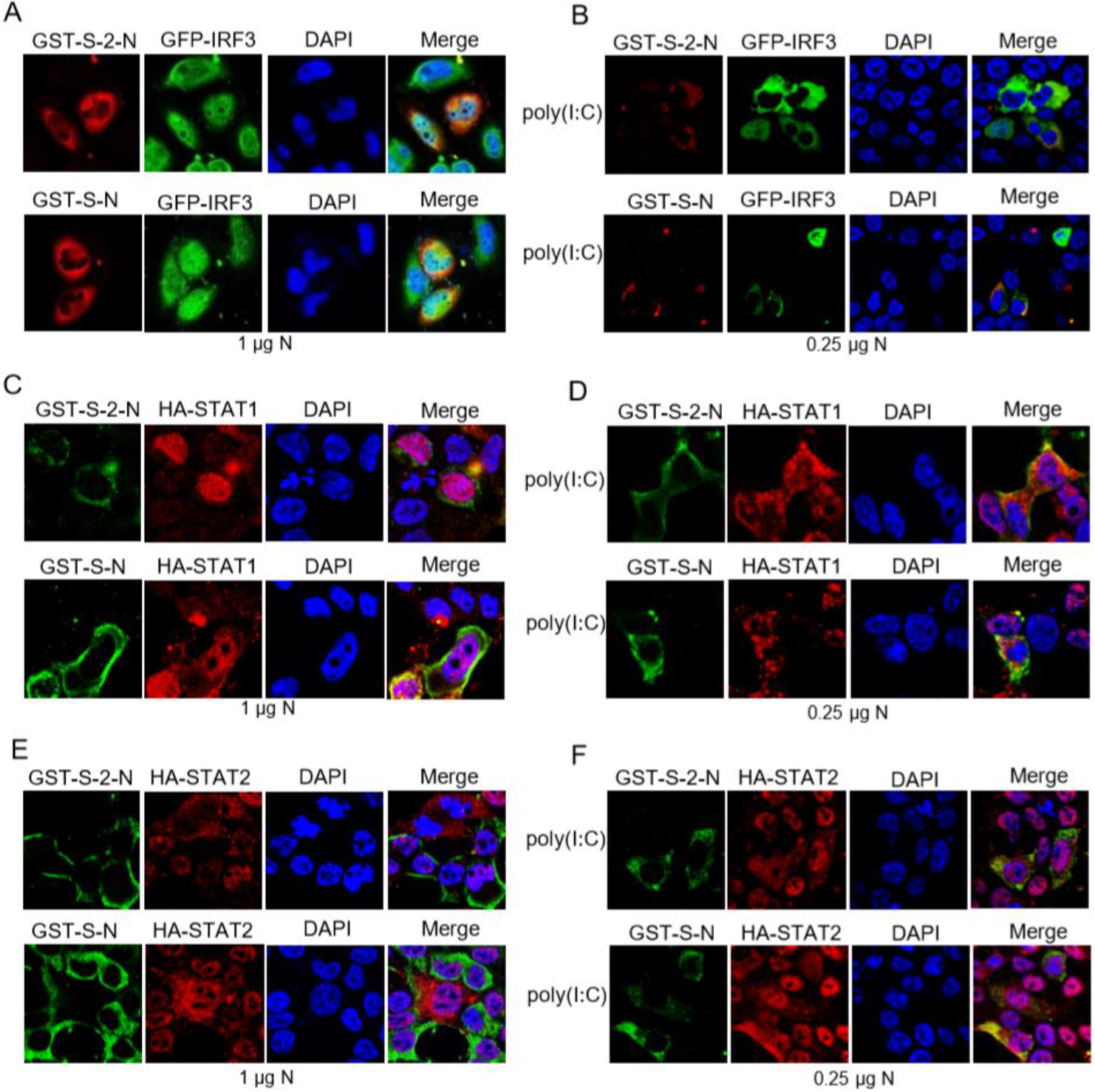
N proteins dually regulate nuclear translocation of IRF3, STAT1 and STAT2. (A) HepG2 cells were co-transfected with GFP-tagged IRF3 and 1 μg GST-tagged SARS-CoV-2 or SARS-CoV N proteins (S-2-N or S-N) plasmids in 24-well plate. At 24 h post-transfection (hpt), cells were subjected to immunofluorescence with anti-GST antibody. (B) HEK293T cells were co-transfected with GFP-tagged IRF3 and 0.25 μg GST-tagged S-2-N or S-N proteins plasmids in 24-well plate, together with poly(I:C). At 24 hpt, cells were subjected to immunofluorescence. Green: IRF3 signal; Red: N protein signal; Blue: DAPI (4, 6-diamino-2-phenyl indole, nuclei staining). Merge indicate the merged red, green, and blue channels. (C, E) HEK293T cells were co-transfected with HA-tagged STAT1 (C) or STAT2 (E) and 1μg GST-tagged N proteins plasmids in 24-well plate. At 24 hpt, cells were subjected to immunofluorescence with anti-GST and HA antibodies. (D, F) HEK293T cells were co-transfected with HA-tagged STAT1 (D) or STAT2 (F) and 0.25μg GST-tagged N proteins plasmids, together with poly(I:C). At 24 hpt, cells were subjected to immunofluorescence. Green: N protein signal; Red: STAT signal; Blue: DAPI. Merge indicate the merged red, green, and blue channels.

### N protein inhibits IFN-I production through TRIM25

The tripartite motif protein 25 (TRIM25) is an E3 ubiquitin ligase that participates in various cellular processes, including regulating the innate immunity against virus infections (18–21). We explored whether the low-dose SARS-CoV-2 N protein inhibits IFN-I signaling through TRIM25, and found that the inhibition of IFN-I activity by SARS-CoV-2 N protein were rescued at both protein and transcriptional levels through overexpression of TRIM25 (Fig.4A-C). We further examined the ability of low-dose SARS-CoV-2 N protein to inhibit IFN-I signaling in TRIM25 knockout cells (sg25). As expected, the SARS-CoV-2 N protein failed to inhibit the ISRE promoter activity and transcriptional activation of IFN-I and ISGs in sg25 cells (Fig. 4D, E). In addition, low-dose SARS-CoV-2 N protein increased the phosphorylation of IRF3 and STAT2 in sg25 cells, indicating that TRIM25 was responsible for the inhibitory effect of SARS-CoV-2 N protein on IFN-I signaling (Fig. 4F). Collectively, these data demonstrated that TRIM25 is involved in the inhibition of IFN-I signaling by low-dose SARS-CoV-2 N protein.

**FIG 4.**
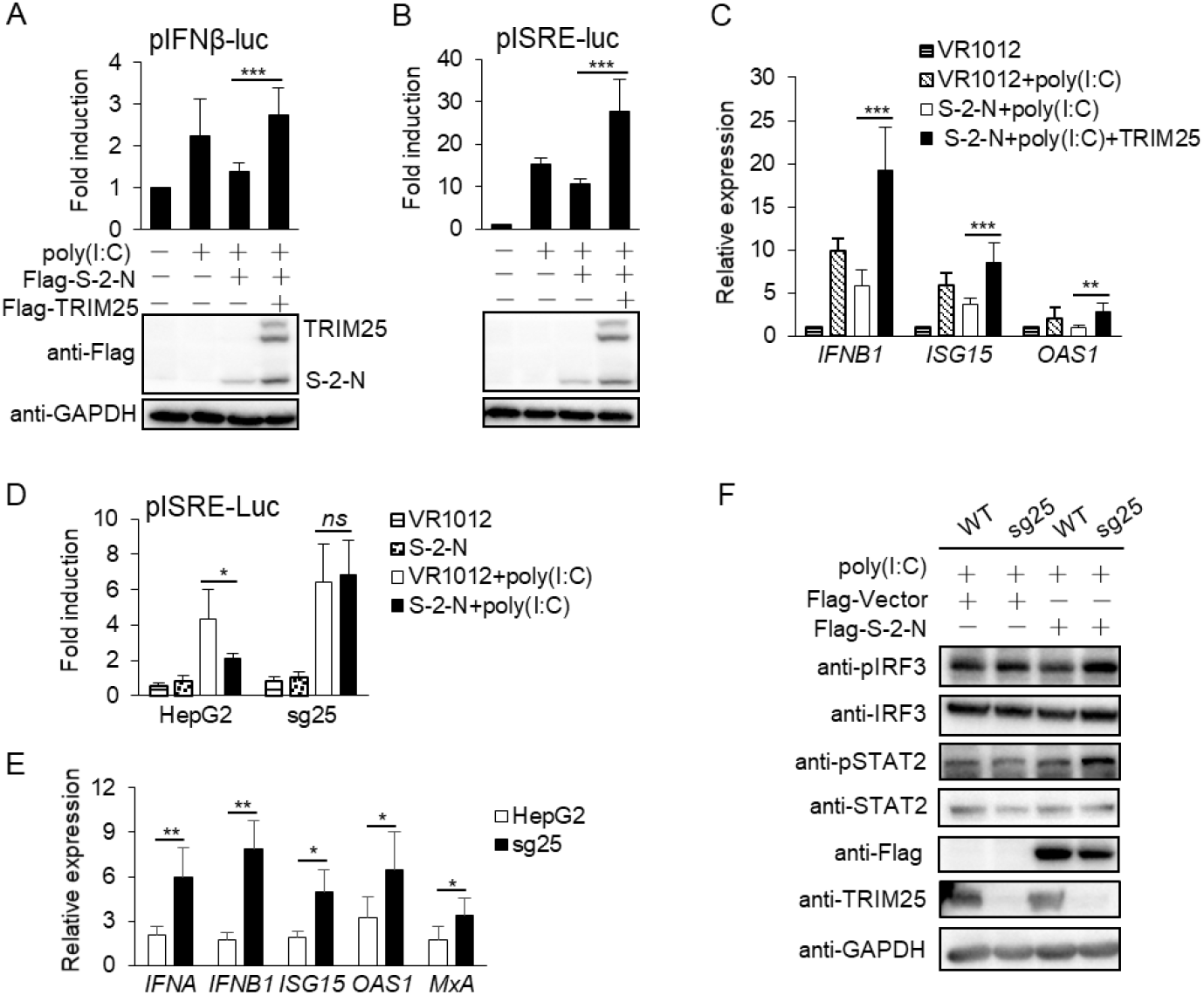
Low-dose N protein inhibits IFN-I production through TRIM25. (A, B) HEK293T cells were co-transfected with reporter plasmid IFN-β (A), or ISRE (B), and the indicated plasmids expressing TRIM25 and 0.25 μg SARS-CoV-2 N (S-2-N) protein, together with or without poly(I:C). At 24 h post-transfection (hpt), cells were harvested and luciferase activity was measured (upper). Expression of S-2-N and TRIM25 detected by Western blot (down). (C) HEK293T cells were transfected with the plasmids expressing 0.25 μg S-2-N alone, or S-2-N and TRIM25, together with or without poly (I:C), VR1012 as an empty control. After 24hpt, the mRNA expression of *IFNB1*, *ISG15* and *OAS1* were determined by qPCR. (D) TRIM25 knockout (sg25) and WT HepG2 cells were transfected with an ISRE reporter and 0.25 μg S-2-N plasmids, together with or without poly(I:C). At 24 hpt, cells were harvested and luciferase activity was measured. (E) sg25 and WT cells were transfected with 0.25 μg S-2-N plasmid and poly(I:C). At 24 hpt, the mRNA levels of *IFNA*, *IFNB1*, *ISG15*, *OAS1* and *MxA* were determined using qPCR. (F) sg25 and WT cells were transfected with 0.25 μg Flag-tagged S-2-N plasmid or empty vector together with poly(I:C). At 24 hpt, the phosphorylation levels of IRF3 and STAT2 were detected by Western blot. Results shown are the mean ± SD of at least three independent experiments. \**P* < 0.05, \*\**P* < 0.01, and \*\*\**P* < 0.001; two-tailed Student’s *t*-test.

### N protein inhibits interaction of TRIM25 and RIG-I

Both SARS-CoV and MERS-CoV N proteins can interfere with the association between TRIM25 and RIG-I by interacting with TRIM25 (22, 23). To determine whether the SARS-CoV-2 N protein specifically binds to TRIM25, we co-transfected SARS-CoV-2 N protein and TRIM25 into HEK293T cells. After 24 h post-transfection, the cell lysates were subjected to immunoprecipitation (IP). We found that TRIM25 was co-immunoprecipitated with SARS-CoV-2 N protein (Fig. 5A), and SARS-CoV-2 N protein was also co-immunoprecipitated with TRIM25 (Fig. 5B). Immunofluorescence revealed that SARS-CoV-2 N protein was co-localized with TRIM25 in the cytoplasm of transfected cells (Fig. 5C). These results indicated that SARS-CoV-2 N protein directly interacts with TRIM25.

**FIG 5.**
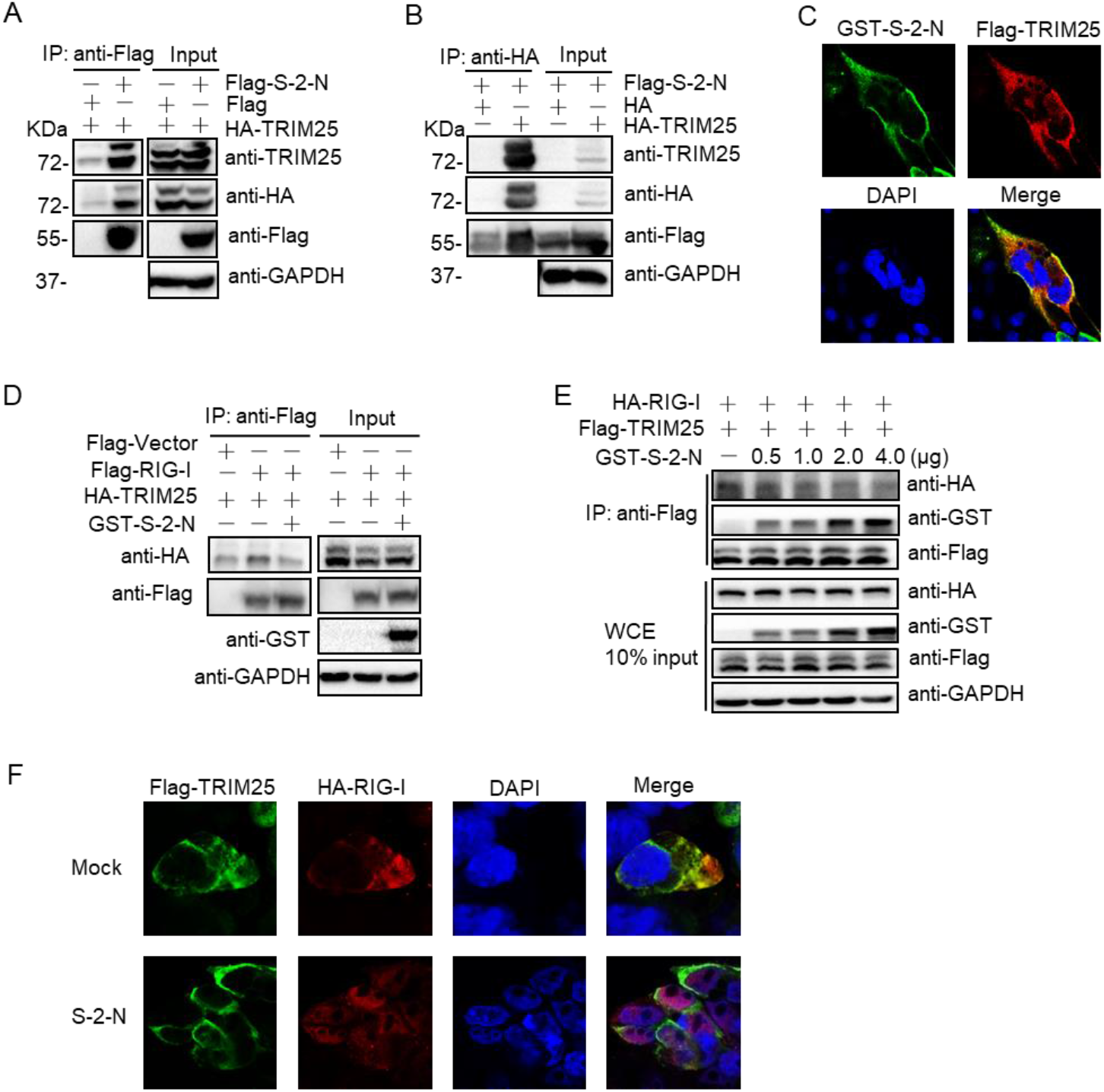
N protein inhibits the interaction of TRIM25 and RIG-I through competitively binding TRIM25. (A) HEK293T cells were transfected with the plasmids expressing Flag-SARS-CoV-2 N (S-2-N) or empty-vector together with HA-TRIM25. After 24 h post-transfection (hpt), the cell lysates were subjected to anti-Flag immunoprecipitation (IP) and analyzed by immunoblot with anti-TRIM25, HA and Flag antibodies, GAPDH was used as a loading control. (B) HEK293T cells were transfected with the indicated plasmids for 24 h, the cell lysates were subjected to anti-HA IP and analyzed by immunoblot. (C) HEK293T cells were transfected with the plasmids expressing GST-S-2-N and Flag-TRIM25. After 24hpt, the cells were subjected to immunofluorescence with anti-GST and Flag antibodies. (D) HEK293T cells were transfected with Flag-RIG-I or empty-vector and HA-TRIM25, together with or without GST-S-2-N plasmids. At 24 hpt, anti-Flag immunoprecipitates were analyzed by immunoblot. (E) HEK293T cells were transfected with Flag-TRIM25 and HA-RIG-I, together with an increasing concentration of GST-S-2-N plasmids for 24h, anti-Flag immunoprecipitates were analyzed by immunoblot. (F) HEK293T cells were transfected with Flag-TRIM25 and HA-RIG-I, together with or without S-2-N plasmids. After 24hpt, the transfected cells were subjected to immunofluorescence with anti-Flag and HA antibodies.

Then, we explored whether SARS-CoV-2 N protein inhibits the interaction of TRIM25 and RIG-I. The plasmids expressing TRIM25 and RIG-I, together with or without SARS-CoV-2 N protein were co-transfected into HEK293T cells, and co-IP assays demonstrated that the interaction ofTRIM25 with RIG-I was inhibited by SARS-CoV-2 N protein in a dose-dependent manner (Fig. 5D, E). Immunofluorescence also revealed that the co-localization of RIG-I and TRIM25 was destroyed by co-expression of SARS-CoV-2 N protein (Fig. 5F). Collectively, these findings demonstrated that SARS-CoV-2 N protein negatively regulates the interaction of TRIM25 with RIG-I.

### N protein inhibits RIG-I ubiquitination through TRIM25

TRIM25 is involved in the K63-linked ubiquitination of RIG-I, which is essential for the stimulation of IFN-I production (24, 25). We next investigated the role of the SARS-CoV-2 N protein in the ubiquitination of RIG-I mediated by TRIM25. First, we examined whether SARS-CoV-2 N protein inhibits RIG-I ubiquitination. HEK293T cells were transfected with RIG-I and ubiquitin plasmids together with SARS-CoV-2 N protein. The results indicated that RIG-I ubiquitination was significantly impaired by SARS-CoV-2 N protein in a dose-dependent manner (Fig. 6A), which was also confirmed by the immunofluorescence analysis (Fig. 6B). We further analyzed whether this ubiquitination inhibitory was mediated by TRIM25, and found that TRIM25 over-expression rescued the suppression of RIG-I ubiquitination caused by SARS-CoV-2 N protein (Fig. 6C). In addition, the RIG-I ubiquitination was reduced in TRIM25-knockout cells, and the inhibition of RIG-I ubiquitination by SARS-CoV-2 N protein was also abolished (Fig. 6D). These results indicated that SARS-CoV-2 N protein inhibits RIG-I ubiquitination through interfering with the interaction of TRIM25 and RIG-I.

**FIG 6.**
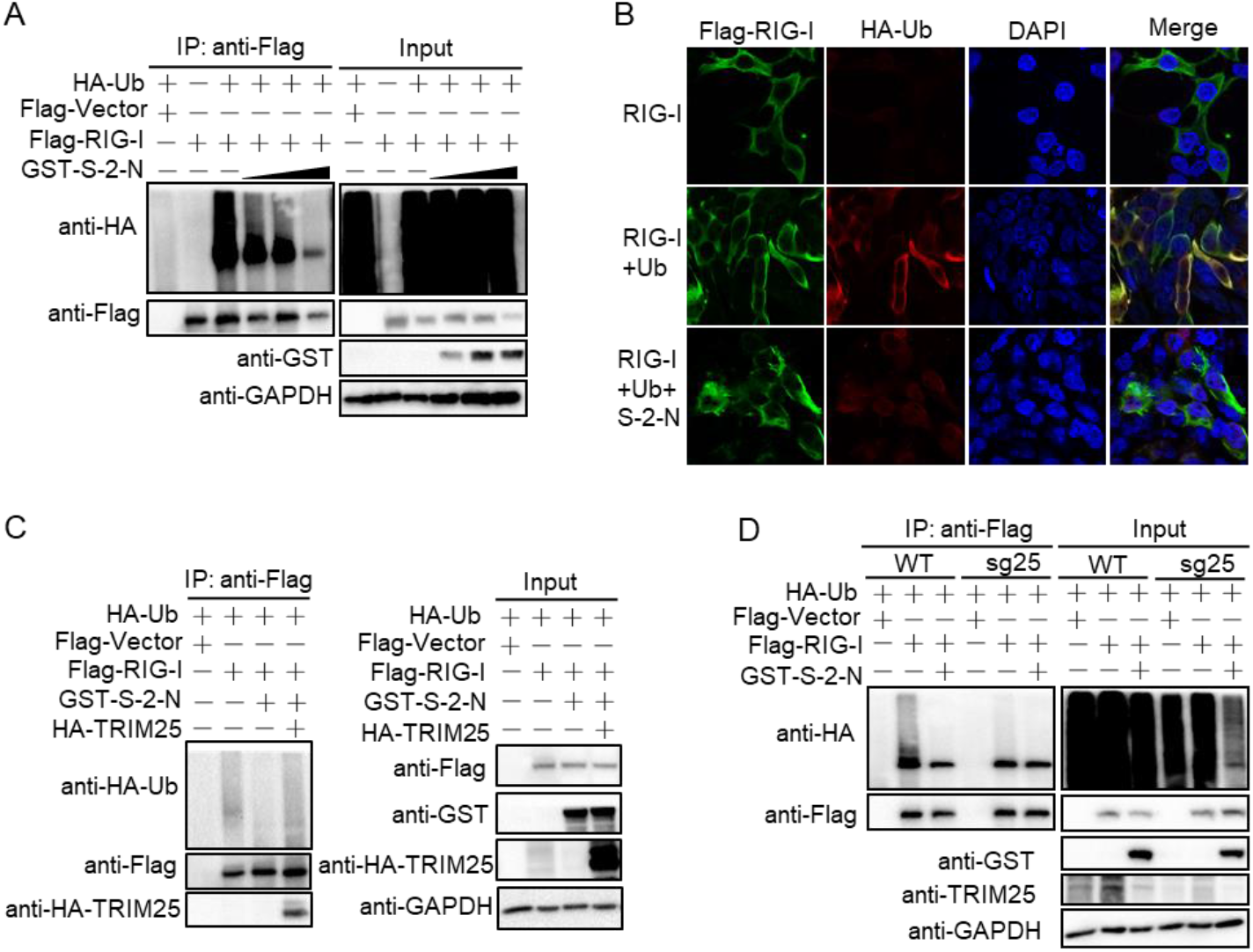
N protein inhibits RIG-I ubiquitination through TRIM25. (A) HEK293T cells were transfected with HA-Ub and Flag-RIG-I plasmids as indicated, together with an increasing GST-SARS-CoV-2 N protein (S-2-N) for 24 h. Anti-Flag immunoprecipitates were analyzed by immunoblot with anti-HA, Flag and GST antibodies. (B) HEK293T cells were transfected with Flag-RIG-I, or Flag-RIG-I and HA-Ub, or Flag-RIG-I, HA-Ub and GST-S-2-N plasmids. At 24 h post-transfection (hpt), the cells were subjected to immunofluorescence analyses with anti-Flag and HA antibodies. (C) HEK293T cells were transfected with the plasmids as indicated, anti-Flag immunoprecipitates were analyzed by immunoblot. (D) sg25 and WT cells were transfected with the indicated plasmids, anti-Flag immunoprecipitates were analyzed by immunoblot.

### Interaction domains of N protein and TRIM25

Specific functional domain of SARS-CoV N is responsible for suppression of IFN-I production (23). We investigated which domain of SARS-CoV-2 N protein is involved in its interaction with TRIM25. Full-length or truncated fragments (aa 1 to 360 and 361 to 419) of SARS-CoV-2 N protein were co-transfected with TRIM25 into 293T cells. We found that both truncated proteins can bind to TRIM25 (Fig. 7A), and were sufficient to suppress IFN-I production (Fig. 7B, C). To determine the key domains for the truncated protein of 1 to 360 aa, we then constructed a series of truncations (Fig. 7F). The protein-protein interactions of these truncations with TRIM25 were examined via co-IP assays, demonstrating that the truncations (aa 1-175 and 252-360) of SARS-CoV-2 N protein can interact with TRIM25, but the truncation of aa 176-251 did not work (Fig. 7D, E).

**FIG 7.**
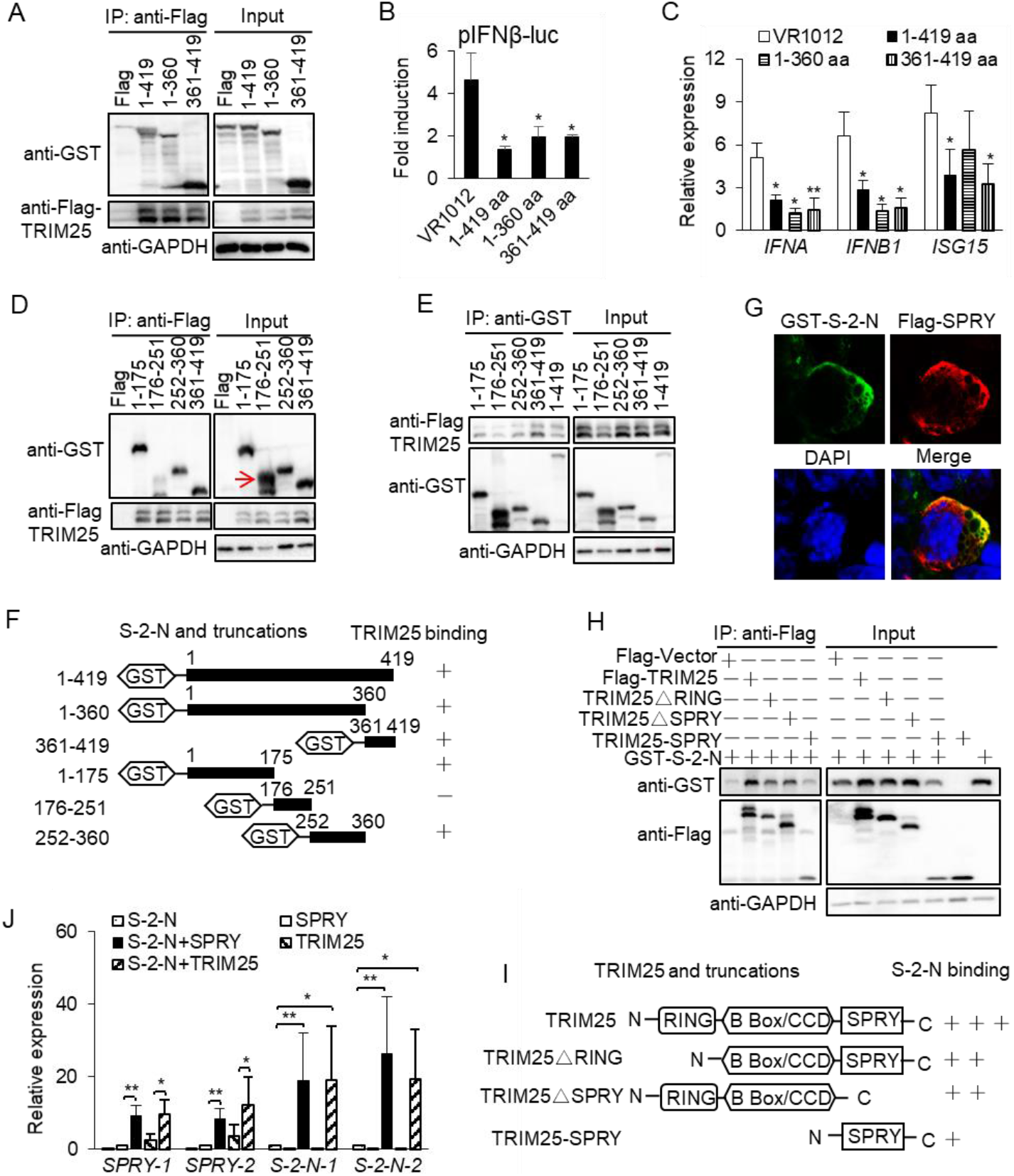
The interaction domains of N protein and TRIM25. (A) HEK293T cells were transfected with the plasmids expressing GST-tagged full-length or truncated SARS-CoV-2 N (S-2-N) protein, together with Flag-TRIM25 or Flag-vector. At 24 h post-transfection (hpt), anti-Flag immunoprecipitates were analyzed by immunoblot with anti-GST and Flag antibodies. (B, C) HEK293T cells were transfected with an IFN-β promoter reporter and the full-length or truncated S-2-N plasmids, then treated with poly (I:C). At 24hpt, luciferase activity was measured (B), and the mRNA expression of *IFNA*, *IFNB1* and *ISG15* were examined using qPCR (C). (D) HEK293T cells were transfected with the indicated GST-tagged truncations of S-2-N and Flag-TRIM25 or Flag-vector. At 24 hpt, anti-Flag immunoprecipitates were analyzed by immunoblot. The arrow indicates the target protein band. (E) HEK293T cells were transfected with the GST-tagged truncations of S-2-N protein and Flag-TRIM25 for 24 h, the cell lysates were incubated using Glutathione Agarose to purify GST-S-2-N truncations. The purified proteins were analyzed by immunoblotting. (F) Domain mapping of the SARS-CoV-2 N protein and TRIM25 association. (G) HEK293T cells were transfected with plasmids expressing GST-S-2-N and Flag-TRIM25-SPRY domain. At 24 hpt, the cells were subjected to immunofluorescence analyses with anti-GST and Flag antibodies. (H) HEK293T cells were transfected with the plasmids expressing Flag-tagged full-length or truncated TRIM25, together with GST-S-2-N protein. At 24 hpt, anti-Flag immunoprecipitates were analyzed by immunoblot. (I) Domain mapping of the association. (J) HEK293T cells were transfected with the indicated plasmids. At 24 hpt, qPCR examined mRNA expression of *SPRY* and *S-2-N* using two pairs primers. Results shown are the mean ± SD of at least three independent experiments. *\*P* < 0.05 and *\*\*P <* 0.01; two-tailed Student’s *t*-test.

As a ubiquitin E3 ligase, TRIM25 includes a RING-finger domain, a SPRY domain, a coiled-coil dimerization domain, and two B-box domains (24). Reciprocally, plasmids expressing SPRY, SPRY deletion and RING-finger deletion truncations were constructed (Fig. 7I). We found that SARS-CoV-2 N protein was co-localized with SPRY domain of TRIM25 in the cytoplasm, which is same as SARS-CoV N protein (23) (Fig. 7G, Fig. S4D). In addition, co-IP results showed that SARS-CoV-2 N protein can interact with both the SPRY- and RING-deleted TRIM25 truncations with the lower binding level than that of full-length TRIM25 (Fig. 7H, Fig. S4A and S4B), indicating that SARS-CoV-2 N protein can interact with the SPRY and other domains of TRIM25, including the RING-finger domain, while only SPRY domain of TRIM25 is responsible for the interaction with SARS-CoV N protein (Fig. S4C-E). These findings clearly defined the interaction domains of the SARS-CoV N protein and TRIM25 (Fig. 7F, I).

We unexpectedly found that the expression of both SARS-CoV-2 N protein and SPRY domain was obviously reduced in the co-transfected cells (Fig. 7H, Fig. S4A, S4B), while the mRNA levels of the SARS-CoV-2 N and SPRY were increased (Fig. 7J). Unlike inhibition of TRIM25-induced IFN-I production, SARS-CoV-2 N protein enhanced the SPRY domain-induced IFN-I production (Fig. S5A), suggesting that SARS-CoV-2 N and SPRY may be degraded by post-transcriptional modifications after the activation of IFN-I signaling. The ubiquitin-mediated degradation of the SARS-CoV-2 N protein and SPRY domain has been excluded in this research (Fig. S5B-D), and the precise mechanism needs further investigation.

## DISCUSSION

COVID-19 patients have undergone a marked immune response, which embodies in the insufficient IFN-I production (immunosuppression) in the early stage of infection and cytokine storm (overactive immune response) in the late stage of infection (6, 26). The two stages of immune reactions are associated with disease severity of COVID-19. Understanding the underlying mechanisms will contribute to development of effective therapies for this emerging coronavirus. In this study, we demonstrated that low-dose SARS-CoV-2 N protein inhibited IFN-I production by disturbing the interaction of TRIM25 and RIG-I and reducing the phosphorylation and nuclear translocation of IRF3, STAT1, and STAT2, whereas high-dose N protein promotes the IFN-I and inflammatory cytokine expression by enhancing the phosphorylation and nuclear translocation of IRF3, STAT1, and STAT2 (Fig. 8).

**FIG 8.**
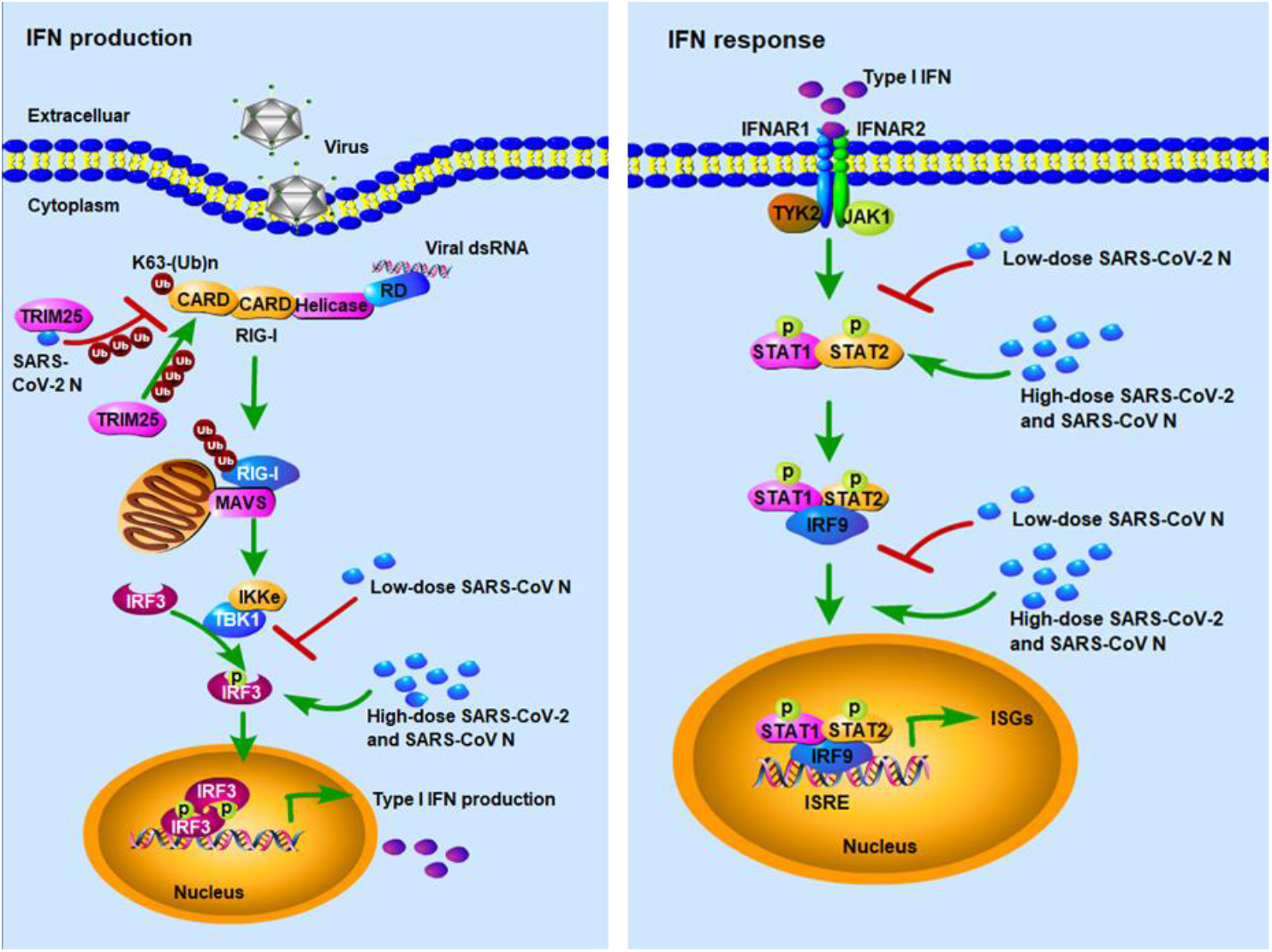
Schematic depiction of the dual regulation role of SARS-CoV-2 nucleocapsid protein against IFN-I signaling. The low-dose SARS-CoV-2 N protein suppresses innate immune response by reducing RIG-I ubiquitination through interacting with TRIM25, and reducing IRF3/STAT1/STAT2 phosphorylation and nuclear translocation; whereas high-dose SARS-CoV-2 and SARS-CoV N proteins promote innate immune response by enhancing the phosphorylation and nuclear translocation of IRF3, STAT1, and STAT2.

Due to the important role of IFN-I in the antiviral immune response, SARS-CoV-2 has employed several different mechanisms to escape from the IFN responses. For example, membrane (M) protein targets MAVS to impair its aggregation (27); NSP1 binds the 40S ribosomal subunit to shutdown mRNA translation of IFNs and ISGs(12); NSP6 binds TANK binding kinase 1 (TBK1) to inhibit IRF3 phosphorylation (28); NSP13 binds TBK1 to block its phosphorylation (28); ORF6 binds importin Karyopherin α 2 to inhibit IRF3 and STAT1 nuclear translocation (10, 11, 28); and ORF3b increases the IFN-antagonistic activity through its truncated C-terminal (29). In addition, SARS-CoV-2 N protein has been suggested to inhibit IFN-I production through interacting with RIG-I and suppressing the phosphorylation and nuclear translocation of STAT1 and STAT2 (16, 17), which supporting our findings that low-dose SARS-CoV-2 N protein could antagonize the IFN-I signaling by inhibiting the phosphorylation and nuclear translocation of IRF3, STAT1, and STAT2. Moreover, our study also showed that low-dose SARS-CoV-2 N protein can inhibit RIG-I ubiquitination through interacting with TRIM25 to suppress IFN-I production (Fig. 8). These mechanistic findings complemented the previous works and detailed the mechanisms of IFN inhibition mediated by SARS-CoV-2 N protein.

SARS-CoV N protein has also shown to inhibit IFN-I production by interfering with TRIM25 (23). However, the domains of N protein and TRIM25 involved in the inhibition and interaction are different in the two coronaviruses. We demonstrated that both N-terminal (aa 1-360) and C-terminal (aa 361-419) of the SARS-CoV-2 N protein could suppress IFN-I production, while another report showed that only the N-terminal (aa 1-361) has inhibitory effect (16). Interestingly, only C-terminal (aa 362-422) of the SARS-CoV N protein is responsible for the suppression of IFN-I production (23). We also found that there are multiple loci in the SARS-CoV-2 N protein, including the N-terminal (aa 1-360 and aa 1-175), C-terminal (aa 361-419) and intermediate region (aa 252-360) could bind to TRIM25, and SARS-CoV-2 N protein could interact with multiple domains of TRIM25 (such as SPRY and RING-finger), so we could conclude that there were multiple loci in both SARS-CoV-2 N protein and TRIM25 involved in their interactions to inhibit IFN-I signaling. On the contrary, only single domain of the SARS-CoV N protein C-terminus (aa 362-422) and TRIM25 SPRY domain was involved in their interaction to inhibit production of IFN-I (23). Theses discrepancies may contribute to the different pathogenicity between SARS-CoV and SARS-CoV-2.

Clinical investigations have shown that the severity of COVID-19 is positively correlated with the cytokine storm, which is characterized by the clinical manifestations of systemic inflammation, hemodynamic instability, hyperferritinemia, and multi-organ failure (30–32). Moreover, both SARS-CoV and MERS-CoV infections can trigger the cytokine storm that contribute to occurrence of ARDS and is the leading cause of death (33, 34). The proinflammatory cytokines IL1B, IL6, IL18, IFNγ, and TNFA are the key components of cytokine storm in the coronavirus infections (35). For example, IL6 is closely correlated with ARDS severity and blood viral load of COVID-19 (36, 37). Our study showed high-dose (1 μg) SARS-CoV-2 N protein can promote expression of IL6, IL1B, and TNFA, demonstrating that SARS-CoV-2 N protein was involved in the cytokine storm of COVID-19.

Low-dose SARS-CoV-2 N protein inhibited RIG-I ubiquitination through interacting with TRIM25 to suppress IFN-I production. In TRIM25 knockout cells, the regulatory effect of high-dose N protein was exactly opposite to that of WT cells (Fig. S6), indicating that high-dose SARS-CoV-2 N protein also activated IFN-I signaling through TRIM25. However, the two-way regulation mechanism of TRIM25 is needed further investigation.

Our results also showed that high-dose SARS-CoV N protein could promote innate immune response by promoting IRF3/STAT1/STAT2 phosphorylation and nuclear translocation. In addition, low-dose SARS-CoV N protein inhibited IFN-I production through TRIM25, whereas high-dose N protein enhanced IFN-I production through TRIM25 (Fig. S6). Considering that both SRAS-CoV and MERS-CoV N proteins can inhibit IFN-I production through interacting with TRIM25(22, 23), we speculate that the N proteins of SARS-like coronaviruses have the common ability to dually regulate innate immune response.

In summary, we demonstrated that the SARS-CoV-2 N protein dually regulates innate immune responses, in which the low-dose N protein suppresses type I interferon (IFN-I) signaling and inflammatory cytokines, whereas high-dose N protein promotes IFN-I signaling and inflammatory cytokines. The SARS-CoV-2 N protein interacts with TRIM25, thereby dually regulating the phosphorylation and nuclear translocation of IRF3, STAT1 and STAT2. Our findings reveal a regulatory mechanism of innate immune responses mediated by the SARS-CoV-2 N protein, which may contribute to development of more effective strategies for controlling COVID-19 and understanding the pathogenesis of SARS-CoV-2 and other SARS-like coronaviruses.

## MATERIALS AND METHODS

### Cells and antibodies

Human embryonic kidney cell HEK293T and hepatic carcinoma cell HepG2 were maintained in Dulbecco’s modified Eagle’s medium (DMEM) (HyClone, Logan, UT) containing 10% inactivated fetal bovine serum (BBI, Shanghai, China), penicillin (100 IU/ml) and streptomycin (100 mg/ml) at 37 °C in a 5% CO_2_ atmosphere. Sg25 cell is derived from HepG2 cell, in which TRIM25 has been knocked out through the CRISPR/Cas9 systems (38).

Antibodies against STAT1, p-STAT1, and STAT2 were purchased from Abcam (Cambridge, MA, USA); anti-IRF3, anti-HA, anti-GST, anti-Flag, anti-GAPDH, CoraLite 594-conjugated IgG and CoraLite 488-conjugated IgG secondary antibodies were obtained from Proteintech (Rosemont, IL, USA); anti-TRIM25 and anti-pIRF3 antibodies were from Cell Signaling technology (Danvers, MA, USA); anti-pSTAT2 antibody was from Sigma-Aldrich (St. Louis, MO, USA).

### Plasmid construction

The IFNβ promoter and ISRE luciferase reporter plasmids, the control reporter plasmid (pGL4.74), and plasmids for Flag- and HA-tagged TRIM25 and ubiquitin were previously described (38, 39). HA-tagged RIG-I plasmid was purchased from Public Protein/Plasmid Library (PPL, Jiangsu, China). GFP-tagged IRF3 and HA-tagged STAT1/2 plasmids were purchased from Miaoling Plasmid Library (Wuhan, China). Flag or GST-tagged RIG-I, SARS-CoV N protein (SARS coronavirus Tor2, NC_004718.3), SARS-CoV-2 N protein (IPBCAMS-WH-01/2019 strain, no. EPI_ISL_402123) and truncations were cloned into VR1012-based expression vector and confirmed by sequencing. The SARS-CoV-2 N protein shares 90.52% aa identity with the SARS-CoV N protein (Fig. S1).

### Transfection and reporter assays

The luciferase reporter assays were performed as previously described (40). Briefly, HEK293T, HepG2 or sg25 cells were transfected with a control plasmid or protein expression plasmids together with the luciferase reporter plasmids using Viafect TM Transfection Reagent (Promega, Madison, WI, USA) or Lipofectamine TM 2000 (Invitrogen, San Diego, CA, USA). After 8 h, the cells were then transfected with poly(I:C) (Invitrogen), 16 h later, promoter activity was measured with Dual-Luciferase® Reporter Assay System (Promega). The relative firefly luciferase activities are normalized to the Renilla luciferase.

### Quantitative real-time PCR (qPCR)

Total RNA was extracted using an EasyPure® RNA Kit (TransGen, Beijing, China), and the first-strand cDNA was synthesized by TransScript First-Strand cDNA Synthesis Super Mix (TransGen). The qPCR was analyzed with SYBR Green Master (Roche, Basel, Switzerland) as previously described (40). The results were normalized by the house-keeping gene GAPDH. Primers used for these analyses are listed in Supplementary Table S1.

### Coimmunoprecipitation and immunoblot analysis

Coimmunoprecipitation and immunoblot analyses were performed as previously described (39). In brief, cells were lysed in lysis buffer (50 mM Tris-HCl, pH 8.0, 150 mM NaCl, 1% NP-40) containing protease and phosphatase inhibitor cocktail (Selleck, Houston, Texas, USA). For coimmunoprecipitation, lysates were incubated overnight with ANTI-FLAG^®^ M2 Affinity Gel (Sigma-Aldrich), EZview™ Red Anti-HA Affinity Gel (Millipore, Billerica, MA, USA), or Pierce Glutathione Agarose (Thermo Scientific, Rockford, IL, USA), then proteins were separated by SDS-PAGE and transferred onto PVDF membranes. After blocking in TBST containing 5% BSA, the blots were probed with primary antibodies. Determination of the band intensities were performed with ChemiDoc XRS^+^ Molecular Imager software (Bio-Rad, Philadelphia, PA, USA).

### Immunofluorescence

HEK293T or HepG2 cells cultured on 12-mm coverslips were transfected with indicated plasmids. After 24 h, cells were fixed with 4% paraformaldehyde, and permeated with 0.5% Triton X-100. After cells were washed with PBST, they were blocked in 1% BSA and stained with primary antibodies, followed by staining with CoraLite 594- or CoraLite 488-conjugated IgG secondary antibodies (41). Nuclei were stained with DAPI (Yesen Biotechnology, Shanghai, China). Fluorescence images were obtained and analyzed using a confocal microscope (FV3000, OLYMPUS).

## Statistical analysis

Data were statistically analyzed by the Student’s *t*-test, and a *P* value of less than 0.05 was considered statistically significant.

## Acknowledgments

This work was supported by grant from National Natural Science Foundation of China (No. 81972873), the Pearl River Talent Plan in Guangdong Province of China (2019CX01N111), and the Science and Technology Innovation Project in Foshan, Guangdong Province, China (2020001000151).

## Author contributions

Q.L., Y.Z., and G.W. designed the experiments. Y.Z., L.S., Z.W., P.W., and W.W. conducted the experiments. G.W., Y.Y., Z.H., G.T., H.H., and S.H. analyzed the data and conducted statistical analysis. Y.Z. wrote the first draft of the paper. All authors contributed to subsequent drafts and approved the final version of the paper.

